# Exploring Size Exclusion Chromatography Columns 20 and 35 nm Pore Size Effect for Isolation of Extracellular Vesicles

**DOI:** 10.1101/2025.10.20.682612

**Authors:** Balazs Kaszala, Vincenzo Scarsella, Kriti Bomb, Jimmy Fay, Barbara Smith, Priyanka Gokulnath, Ridhdhi Desai, Rohit N. Kulkarni, Saumya Das, Marta Garcia-Contreras

## Abstract

Extracellular vesicles (EVs) have shown great promise as minimally invasive biomarkers for a variety of diseases. However, challenges persist regarding EV isolation, particularly in their co-isolation with impurities such as soluble proteins and lipoproteins. Among the methods available for EV isolation, size-exclusion chromatography (SEC) is widely used, as it is reproducible and amenable to high-throughput with a rapid turnaround time. However, its size-based separation leads to the co-isolation of EVs with impurities of similar size. This study, for the first time to our knowledge, compares SEC columns with different pore sizes, 20 and 35 nm, to evaluate their efficacy in non-EV contaminant removal and EV recovery from pancreatic EndoC-ßH1 cell culture media and human plasma. To assess EV purity and yield, we compare EV particle concentration, the presence of unintended co-isolates, and RNA EV cargo. This study demonstrates that smaller pore size SEC columns enhance EV yield and purity, making them ideal for biomarker studies involving limited biological samples or downstream analysis sensitive to contaminants.

## Introduction

Extracellular vesicles (EVs) are small lipid-bilayer membranous particles that carry proteins, nucleic acids, and lipids^1,2^. They are found in most biological fluids, making them a potentially minimally invasive source of biomarkers for various diseases such as cancer, cardiometabolic diseases, neurological diseases or diabetes^3–6^. Furthermore, EVs can transfer their cargo by mediating cell-to-cell communication and have emerged as therapeutic delivery vehicles^7,8^.

Despite advances in the field, there is a lack of standardization and reproducibility between studies, in part due to variable levels of other nanoparticles in EV samples. In cell culture, fetal bovine serum (FBS) is a source of particle and non-particle contaminants^9–11^. While albumin can be used as a replacement for FBS, the bovine serum from which albumin is derived contains secreted vesicles that are considered contaminants in EV preparations^12^. In human plasma, lipoproteins such as APOA and proteins like albumin are also considered contaminants^13^. Moreover, these contaminants have been shown to confound downstream analyses and the quality of the EV preparations^14^. For example, low-density lipoproteins have been shown to interfere with EV analysis, such as tunable resistive pulse sensing in human plasma^13^. These co-isolates may also contain amounts of cargo (e.g. RNA) that are likely to be a source of variability in clinical samples.

Because EVs are less abundant than many of these contaminants, enriching for EVs without co-isolating these contaminants remains challenging. Several methods are currently used for EV isolation, including filtration, polymer-based precipitation, size-exclusion chromatography, differential centrifugation, density gradient, asymmetric flow field-flow fractionation, or immunoprecipitation^15,16^. These methods differ in the EV purity and yield as well as the need for specialized equipment (such as ultracentrifuges) and the time needed to complete the isolation. A commonly used method for isolating EVs is size-exclusion chromatography (SEC), which separates particles based on size, and can isolate intact and functionally active EVs^17^. SEC columns can present different pore sizes in the resin used to fill the columns, and the column pore size is crucial for effective eparation^18^. Larger molecules are excluded from the pores and elute faster in comparison to smaller molecules that penetrate the pores and elute later. For EV separation, the ideal pore size should range from 20–30 nm to 150–200 nm, corresponding to the size of the smallest and largest EVs respectively. However, SEC may also co-isolate contaminants that fall within the size range as EVs^19–21^. For this reason, to minimize confounding effects on downstream applications of the study, it is important to assess the presence of contaminants in the EV preparations.

Here, we comprehensively compare multiple SEC columns with different pore size to characterize the absence of contaminants as well as the EV recovery and yield, in a cell culture model of pancreatic EndoC-βH1 cells^22–24^ (where the cell culture media contains bovine serum albumin as a contaminant), as well as human plasma containing contaminants such as albumin and lipoproteins. To the best of our knowledge, this is the first study comparing SEC columns with 20 and 35 nm pore sizes for EV isolation and contaminant removal.

## Materials and methods

### Cell Culture

We used EndoC-βH1 cells, a widely used human β-cell line routinely used for *in vitro* studies in islet biology^25–27^. EndoC-βH1 cells were cultured in DMEM low glucose (1g/L) (Catalog number 11885084, Gibco, Thermo Fisher Scientific), 2% BSA fraction V (Catalog number P-753-100G, Boston BioProducts), 50 µM 2-mercaptoethanol (Catalog number M6250-100ML, Sigma-Aldrich), 10 mM nicotinamide (Catalog number N0636-100G, Sigma-Aldrich), 5.5 µg/mL transferrin (Catalog number 10652202001, Sigma-Aldrich), 6.7 ng/mL sodium selenite (Catalog number 214485-5G, Sigma-Aldrich), and 1% penicillin and streptomycin (Catalog number 15140122, Gibco, Thermo Fisher Scientific). The cell culture plates were precoated with 1% ECM (Catalog number E1270, Sigma-Aldrich) and 2 µg/ml fibronectin from bovine plasma (Catalog number F1141, Sigma-Aldrich). The cells were passaged 1:2 every week, cultured at 37 °C with 5% CO_2_, and tested for mycoplasma contamination using MycoAlert PLUS Mycoplasma Detection Kit (Catalog number LT07-218, Lonza).

### Human plasma samples

Human pooled plasma was commercially obtained (Catalog number CCN-10, Biologic) and reported to contain platelet-poor plasma from at least 20 healthy donors aged between 18 and 66.

### Extracellular Vesicle Isolation by Size Exclusion Chromatography

Before EV isolations, conditioned EndoC-βH1 cell culture media was collected and centrifuged at 500 g for 5 minutes to remove cells, then at 2,000 g for 10 minutes to remove dead cells and debris. Media was then filtered through a 0.80 µm and further concentrated using a 100KD filter (Catalog number UFC710008, Millipore). For the human plasma, we centrifuged it at 2,000 g for 10 minutes to remove debris. For both types of samples, the SEC columns were used as follows. The Izon qEV35 and qEV20 SEC columns with pore size of 35 nm and 20 nm respectively (Catalog number ICO-35 and Catalog number ICO-20, Izon) were used with the Automatic Fraction Collector (AFC) V2 from Izon (Catalog number AFC-V2, Izon) with default instrument settings. Briefly, columns were washed with PBS three times manually, and after 500 µL of EndoC-βH1 concentrated conditioned media or 500 µL of human plasma were loaded into the column, six fractions of 500 µL each were collected with instrument default methods. For EV Isolations, fractions 1 to 4 were combined and concentrated to 250 µL using a 10KD filter (Catalog number UFC801008, Millipore). APEX6B (Catalog number Apex 6B SEC Columns, Everest Biolabs) and APEX4B (Catalog number Apex 4B SEC Columns, Everest Biolabs) SEC columns with 20 nm and 35 nm pore size were used with their corresponding instrument, an Ascent Instrument from Everest Biolabs. Briefly, columns were washed with PBS with instrument default settings, and after 500 µL of EndoC-βH1 concentrated conditioned media or 500 µL of human plasma were loaded into the column, six fractions of 500 µL each were collected. For EV Isolations, fractions 2 to 4 were combined and concentrated to 250 µL using a 10KD filter (Catalog number UFC801008, Millipore).

### Microfluidic resistive pulse sensing

Microfluidics resistive pulse sensing measurements were performed with the nCS1 instrument (Spectradyne, Torrance, CA). Isolated extracellular vesicle samples were diluted 1:100 in 1% Tween 20 in 1× PBS (PBST) and loaded onto polydimethylsiloxane cartridges (diameter range 65 nm to 400 nm). A different cartridge was used for each sample and replicate. Approximately 10 µL of the diluted sample (1:100) was used, and events were recorded for each sample. Results were analyzed using the nCS1 Data Analyzer (Spectradyne, Torrance, CA).

### Transmission Electron microscopy

Isolated extracellular vesicles were imaged by TEM. Briefly, 10 μL of each sample was freshly thawed and adsorbed to glow-discharged carbon-coated 400 mesh copper grids by flotation for 2 min. Three consecutive drops of 1× Tris-buffered saline was prepared on Parafilm. Grids were washed by moving from one drop to another, with a flotation time of 10 seconds on each drop. The rinsed grids were then negatively stained with 1% uranyl acetate (UAT) with tylose (1% UAT in deionized water (dIH_2_O), double-filtered through a 0.22-μm filter). Grids were blotted, and then excess UAT was aspirated, leaving a thin layer of stain. Grids were imaged on a Hitachi 7600 TEM operating at 80 kV with an XR80 charge-coupled device (8 megapixels, AMT Imaging, Woburn, MA, USA).

### Immunoblotting

Protein samples were measured by BCA assay (Catalog number 23227, Thermo Fisher). For the fractions equal volume was loaded, while for the rest of the Western blots equal amount of protein was loaded per well. The pellets were lysed in 10X RIPA buffer (Catalog number #9806, Cell Signaling) and protease inhibitors (Catalog number 78425, Thermo Fisher) for 15 min on ice. Samples were spun down for 15 min at 14,000 rpm in a tabletop centrifuge at 4°C. Fractions or lysates were mixed with 4× TGX sample buffer (Catalog number 1610747, Bio-Rad) under non-reducing conditions (for EV tetraspanin blotting) or under reducing conditions, boiled for 5 min at 95°C, and run into a PAGE gel electrophoresis on a 4%–20% Criterion TGX Stain-Free Precast gel (Catalog number 5678094, Bio-Rad). Proteins were transferred to a polyvinylidene fluoride (PVDF) membrane using the turbo transfer system (Bio-Rad) using the mixed molecular weight program. After membranes were blocked for 1 h in 5% BSA in TBS + 0.05% Tween-20 (TBS-T), blots were incubated overnight at 4°C with the indicated primary antibodies in blocking buffer (Table 1). Then blots were washed 3 times with TBS-T and incubated for 1 h at room temperature with secondary antibodies (Table 1). Finally, membranes were washed 3 times with TBS-T and developed with SuperSignal West Femto (Catalog number 34095, Thermo Fisher) on an iBright FL1500 (Thermo Fisher) in chemiluminescence mode.

**Table 1:** List of antibodies.

| Antibody | Catalog Number | Manufacturer | Dilution |
| --- | --- | --- | --- |
| **Western blot** |  |  |  |
| Bovine Serum Albumin Polyclonal Antibody | A11133 | Thermo Scientific | 1:1000 |
| Albumin Antibody | 4929S | Cell Signaling | 1:1000 |
| Anti-ApoB | NB200-527 | Novus | 1:1000 |
| Anti-ALIX antibody | ab88388 | Abcam | 1:1000 |
| Anti-GM130 | MA5-35107 | Thermo Scientific | 1:1000 |
| CD63 Mouse anti Human, Unlabeled, Clone: H5C6, BD | 556019 | BD | 1:1000 |
| Anti-rabbit IgG, HRP-linked Antibody | 7074S | Cell signaling | 1:5000 |
| Anti-mouse IgG, HRP-linked Antibody | 7076S | Cell signaling | 1:5000 |
| **Nanoflow Cytometry** |  |  |  |
| APC anti-human CD9 Antibody | 312108 | BioLegend | undiluted |
| APC anti-human CD81 (TAPA-1) Antibody | 349509 | BioLegend | 1:20 |
| APC anti-human CD63 Antibody | 353008 | BioLegend | 1:20 |

### Human Tetraspanin ELISA

The human tetraspanin ELISA (Catalog number Atlas Human EV ELISA Kit, Everest Biolabs), was run as per the manufacturer’s instructions. Briefly, EV standards and samples were diluted (1:60 for human plasma EVs, 1:6 for EndoC-ßH1 EVs) in sample diluent buffer and the antibody cocktail was added at 1:1 ratio, then incubated overnight at 4°C. Next plates were washed with 1x wash buffer five times and incubated with TMB substrate solution for 10 minutes in the dark on a plate shaker at 600 rpm. Then the stop solution was added to each well and subjected to shaking for 1 minute at 300 rpm. Absorbance was measured at 450 nm using a plate reader (SpectraMax® iD3, Molecular Devices). All samples were run in two technical replicates and three biological replicates.

### APOA ELISA

Atlas APOA ELISA Kit (Catalog number Atlas APOA ELISA Kit, Everest Biolabs) was run as per the manufacturer’s instructions. Briefly, EV standards and samples were diluted 1:60 in sample diluent buffer and the antibody cocktail was added at 1:1 ratio, then incubated overnight at 4°C. Next, plates were washed with 1x wash buffer five times and incubated with TMB substrate solution for 10 minutes in the dark on a plate shaker at 600 rpm. Then the stop solution was added to each well and placed on shaker for 1 minute at 300 rpm. Absorbance was measured at 450 nm using a plate reader (SpectraMax® iD3, Molecular Devices). All samples were run in two technical replicates and three biological replicates.

### Nanoflow cytometry of EV surface markers

Isolated EVs were diluted in 0.01 M HEPES buffer to a particle concentration of approximately 1× 10^10^ particles/ml for staining. Briefly 9 µL of the sample was incubated with 1 µL of the corresponding antibodies previously diluted (1:20) in the dark for 30 minutes (Table 1). Then sample was diluted 100-fold prior to analysis in a nanoflow cytometer (Flow Nanoanalyzer, NanoFCM). Before analysis, the instrument was calibrated for particle concentration and for size distribution NanoFCM™ Silica Nanospheres Cocktail #1 with different sizes 68 to 155 nm (Catalog number S16M-Exo, NanoFCM). Samples were run and data collected for 1 minute with a pressure of 1.0 kPa. Data was analyzed using data were analyzed by the NanoFCM Software V1.17 (NanoFCM).

### qRT-PCR

RNA isolation was performed using miRNeasy® Mini Kit (Catalog number 217004, Qiagen) as per manufacturers protocol. Reverse transcription was performed using miRCURY® LNA® RT Kit (Catalog number 339340, Qiagen) according to the manufacturer’s instructions. RNA integrity was determined with Nanodrop (Nanodrop One, Thermo Fisher Scientific) and a 4150 TapeStation System (Agilent Technologies, Santa Clara, CA, USA). Quantitative real-time PCR (qRT-PCR) was performed using miRCURY® LNA® SYBR Green PCR Kit (Catalog number 339346, Qiagen) and the QuantStudio 6 Flex instrument (Applied Biosystems). Primers for hsa-miR-21-5p (Catalog number 339306/GeneGlobe ID YP00204230), hsa-miR-30d-5p (Catalog number 339306/GeneGlobe ID YP00204032), and the C. elegans miR-39 spike-in control (Catalog number 339306/GeneGlobe ID YP00203952) were purchased from Qiagen. Relative miRNA levels from EV isolates were established against C. elegans miR-39 mimic spike-in control, using the ΔΔCq method.

### Statistical Analysis

Statistical analysis was performed using GraphPad Prism 10 software (version 10.3.1, GraphPad). The student’s t-test and one-way ANOVA with multiple comparisons was used for statistical comparison between the columns, with *p* values of 0.05 or less considered significant (\**p*<0.05, \*\**p*<0.01, \*\*\**p*<0.001, \*\*\*\**p*<0.0001).

## Results

### Characterization of EndoC-βH1 Conditioned Cell Culture Media Isolated Individual Fractions by Size Exclusion Chromatography

EndoC-βH1 cells were grown in media containing bovine serum albumin (BSA), which is considered a contaminant and can compromise the purity of EV preparations. To assess the presence of BSA in the fractions (F) isolated from the EndoC-βH1 conditioned media using size exclusion chromatography (SEC) columns with different sized pores (35 or 20 nm) from two different manufacturers (Figure 1A), we performed Western blot analysis (Figure 1B). In both APEX columns (20 nm and 35 nm), we detected BSA primarily in fractions 3 to 6. In the qEV35 column (35 nm), we detected the presence of BSA primarily in fractions 3 to 6, while in the qEV20 column (20 nm), we detected the presence of BSA in all fractions. We quantified the BSA signal intensity per fraction from triplicate samples and normalized to the BSA content of F6. For APEX columns, no significant differences were observed in the distribution of BSA across fractions (Figure 1C), whereas for the IZON columns, the qEV20 column showed a higher proportion of BSA in most of the fractions (Figure 1D). We next determined the number of particles (including EVs) per fraction using a tetraspanin ELISA (Figure 1E-F). For the APEX6B column, we found fractions F3 and F4 to have the highest particle concentration (5.71×10^10^±9.91×10^9^ and 3.37×10^10^±1.18×10^10^ particles/mL, respectively), while for APEX4B F4 and F5 exhibited the greatest particle concentrations (7.39×10^10^±3.43×10^9^ particles/mL and 7.65×10^10^±2.17×10^9^ particles/mL). For the APEX columns, F2-4 are the fractions where the EVs are expected to be eluted. For the qEV20 column, we found F1 and F2 to have the highest particle concentrations (7.90×10^10^±1.96×10^8^ and 7.89×10^10^±1.51×10^8^ particles/mL), which was also observed for the qEV35 column (2.82×10^10^±1.02×10^10^ and 2.25×10^10^±1.57×10^9^ particles/mL, respectively). For these columns, F1-4 are the expected fractions for elution of EVs. When comparing the fractions for both columns, F1 and F2 have higher particle counts in the qEV35 and qEV20 fractions, while F3 and F4 have higher particle counts in the APEX6B and APEX4B fractions. F5 and F6 have similar amounts of particles for all columns except for APEX4B, although these fractions are not used for downstream EV analysis.

**Figure 1:**
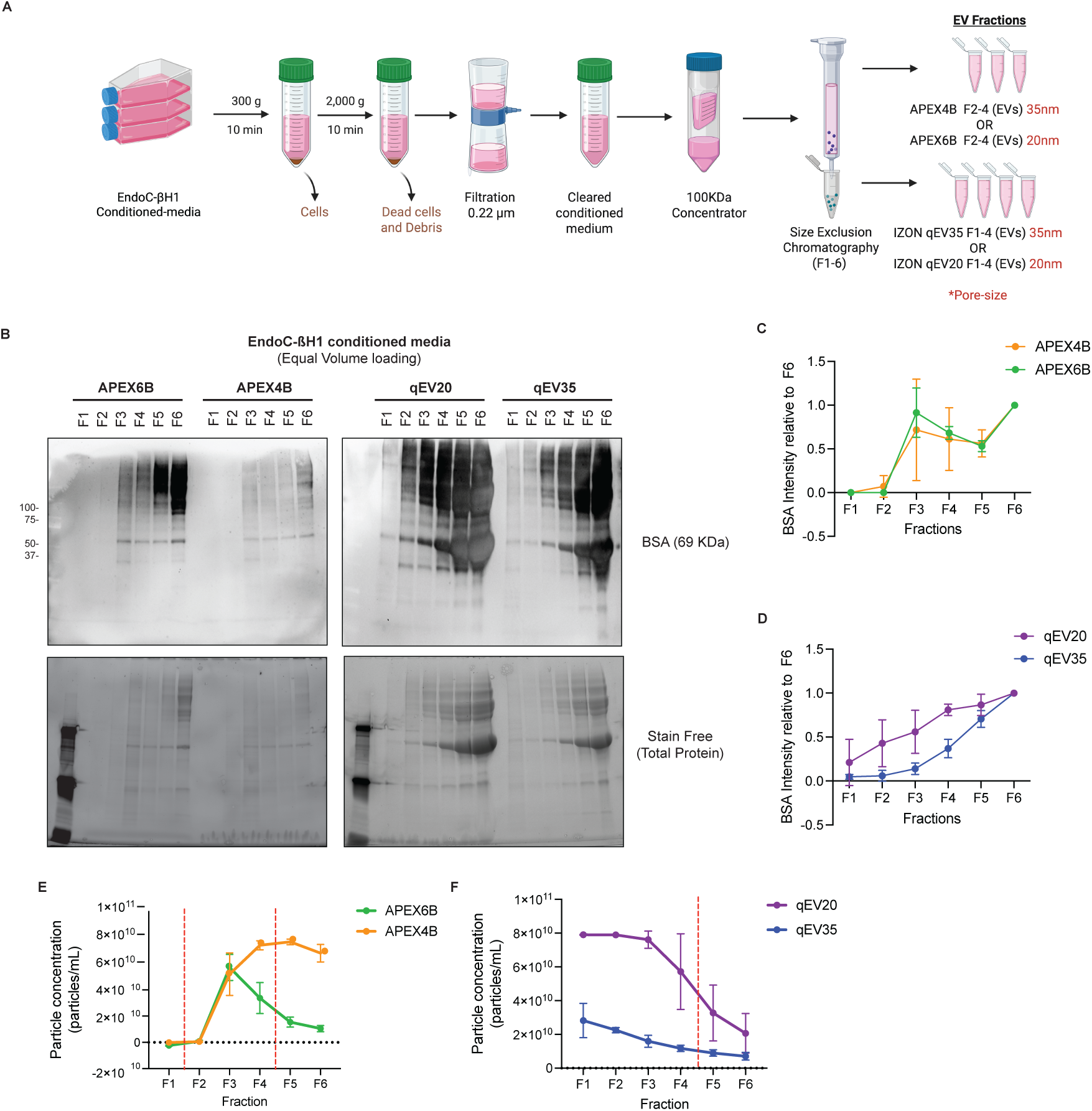
Comparison of isolated fractions from EndoC-βH1conditioned media using different SEC columns. **A**. Schematic Illustration of Isolation workflow. **B**. Representative Western blot of media contaminant Bovine Serum Albumin (BSA), and relative intensity of BSA to F6 (N=3). **C**-**D**. Quantification of human serum albumin present on the western blot fractions of EndoC-βH1 EVs (*n*=3 per group). **E-F**. Particle concentration in the isolated fraction from both SEC columns quantified by TSPAN ELISA (*n*=3) (Data are mean particles/ml ± SD). The red dotted line separates the fractions used for EV isolations – F2-4 for APEX and F1-4 for IZON.

### Comparison of Isolated EVs from EndoC-βH1 Conditioned Cell Culture Media

To investigate the yield and purity of each isolation, we combined and concentrated F2-4 for APEX columns and F1-4 for IZON columns following each manufacturer’s protocol. Morphologically, EVs isolated from both pore sizes looked very similar by TEM, with both showing the typical cup-shaped morphology (Figure 2A). Particle size and concentration were measured by Microfluidic Resistive Pulse Sensing (MRPS) and nanoflow cytometry. For MRPS, all columns showed that isolated EVs had a diameter of 60-80 nm (Figure 2B). The particle concentration for the APEX6B column yielded 1.17×10^11^±1.53×10^10^ particles/mL and 4.60×10^10^±7.21×10^9^ particles/mL for APEX4B, while the qEV20 column yielded 5.17×10^11^±2.87×10^11^ particles/mL and qEV35 column yielded 1.90×10^11^±3.00×10^10^ particles/mL. In general, the columns with 20 nm pore size yielded significantly higher particle concentrations. When analyzed by nanoflow cytometry, mean EV diameter was 77.3±19.7 nm for the APEX6B column, 73.0±22.7 nm for APEX4B, 90.5±19.1 nm for qEV20 and 79.9±18.9 nm for qEV35 (Table 2). Furthermore, the average measured particle concentrations were 1.38×10^10^±3.38×10^8^ particles/mL for APEX 6B, 9.33×10^9^±3.80×10^9^ particles/mL for APEX4B, 1.58×10^10^±6.91×10^8^ particles/mL for qEV 20, and 1.80×10^10^±8.37×10^8^ particles/mL for the qEV35 column (Table 3). To further characterize the EVs, we used immunoblotting to check for the presence of commonly used EV internal and membrane markers (Alix and CD63, respectively) and a negative control (the Golgi apparatus marker GM130) (Figure 1C), as recommended by the Minimal information for studies of extracellular vesicles (MISEV) guidelines^28^. Columns with 35 nm pore size showed significantly higher expression of Alix, and the APEX6B column was shown to have significantly increased expression of CD63 over the APEX4B column. GM130 was not detected in EVs from any column (Figure 2D). Additionally, we calculated EV particle concentrations using a tetraspanin ELISA by measuring the total concentration of the EV fractions for each column. Here we found qEV20 samples to have a significantly higher particle concentration than qEV35 and no significant difference in the APEX samples (Figure 2E). We also measured the total protein concentration of pooled EV samples isolated from both columns using the BCA assay. The protein detected here includes the EV protein as well also any protein contaminants that may have been present in the sample; we found samples isolated with columns with 20 nm of pore size had a higher total protein concentration (Figure 2F).To characterize the relative sample purity, we calculated the ratio of EV particle-to-protein concentration ratio as previously described^29^. We found that APEX4B and qEV35 samples have a significantly higher particle/protein ratio than APEX6B and qEV20 samples (Figure 2G). Finally, to further characterize surface EV markers - the tetraspanins CD9, CD63, and CD81 - we performed nanoflow cytometry (Figure 2H). For APEX6B 54.4±0.95% and for APEX4B 44.7±2.01% of EVs were positive for CD9, while 51.4±0.95% of qEV20 EVs and 51.9±1.36% of qEV35 EVs were positive for CD9 (Table 4). For APEX6B 6.73±0.84% and for APEX4B 13.7±3.27% of EVs were positive for CD63, while 12.6±2.00% of qEV20 EVs and 11.0±1.61% of qEV35 EVs were positive for CD63 (Table 4). For APEX6B 61.7±0.67% and for APEX4B 52.8±4.90% of EVs were positive for CD81, while 44.2±1.01% of qEV20 EVs and 50.0±1.15% of qEV35 EVs were positive for CD81 (Table 4). CD9 and CD81 expression were higher in APEX6B EVs in comparison to APEX4B EVs, while APEX4B EVs displayed increased expression of CD63 compared to APEX6B EVs. qEV35 EVs showed higher CD81 expression than qEV20 EVs.

**Figure 2:**
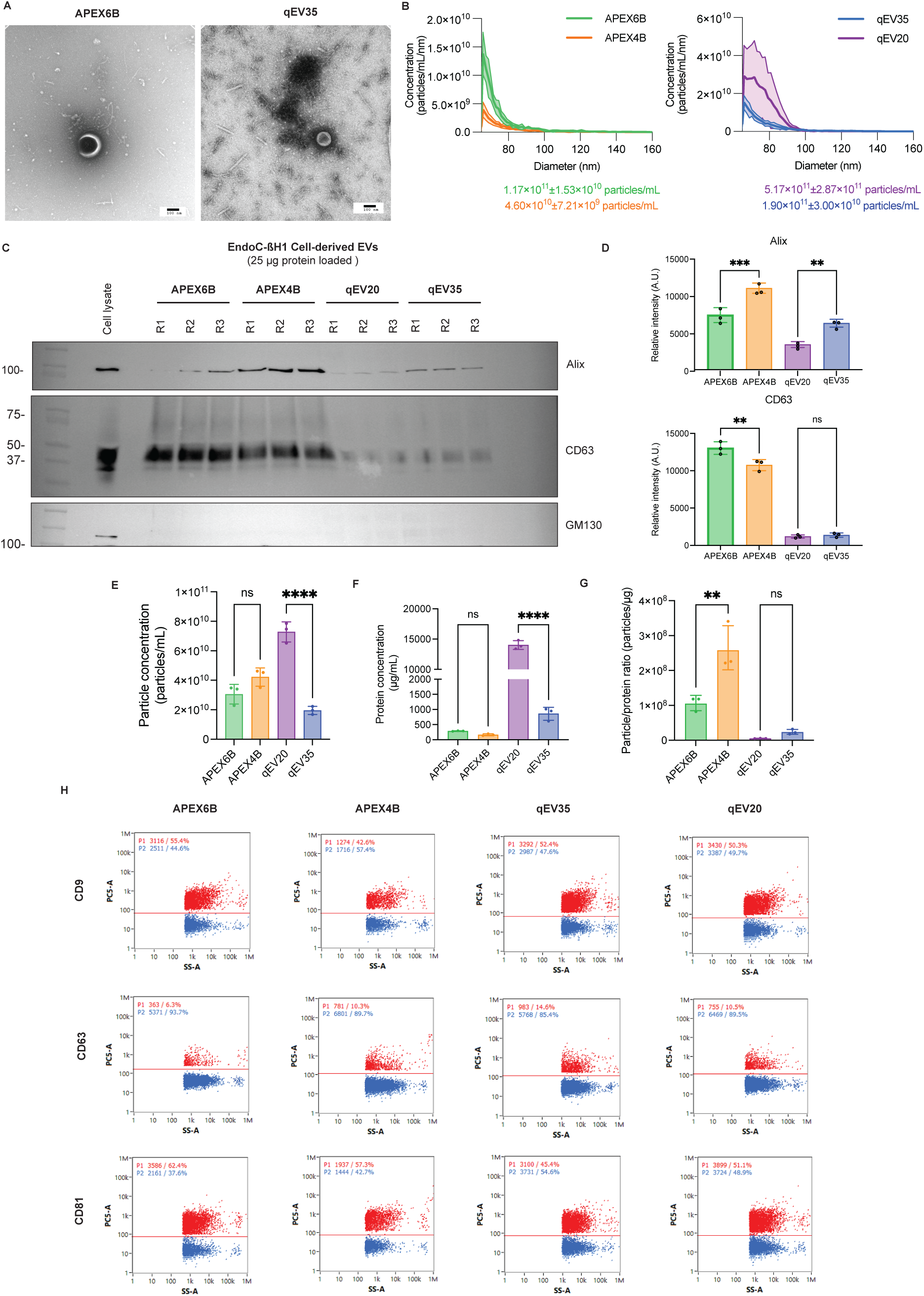
Characterization of EVs isolated from EndoC-βH1 conditioned media by different SEC columns. **A**. Representative images captured by Transmission electron microscopy, Scale=100nm. **B**. Size distribution measured by microfluidic resistive pulse sensing represented as mean± SD (*n*=3).**C-D**. Representative Western blot and quantification of EV markers of EndoC-βH1 EVs (*n*=3). **E**. Particle concentration of the isolated EVs (*n*=3). **F**. EndoC-βH1 EVs protein concentration by BCA (*n*=3). **G**. Particle concentration to protein ratio of isolated EVs (*n*=3) **H**. Representative flow cytometry plots of the detection of EV surface markers CD9, CD63, or CD81 on isolated EVs. All data are means ± SD and *n*=3. \**p*<0.05, \*\**p*<0.01. R=replicate.

**Table 2:**
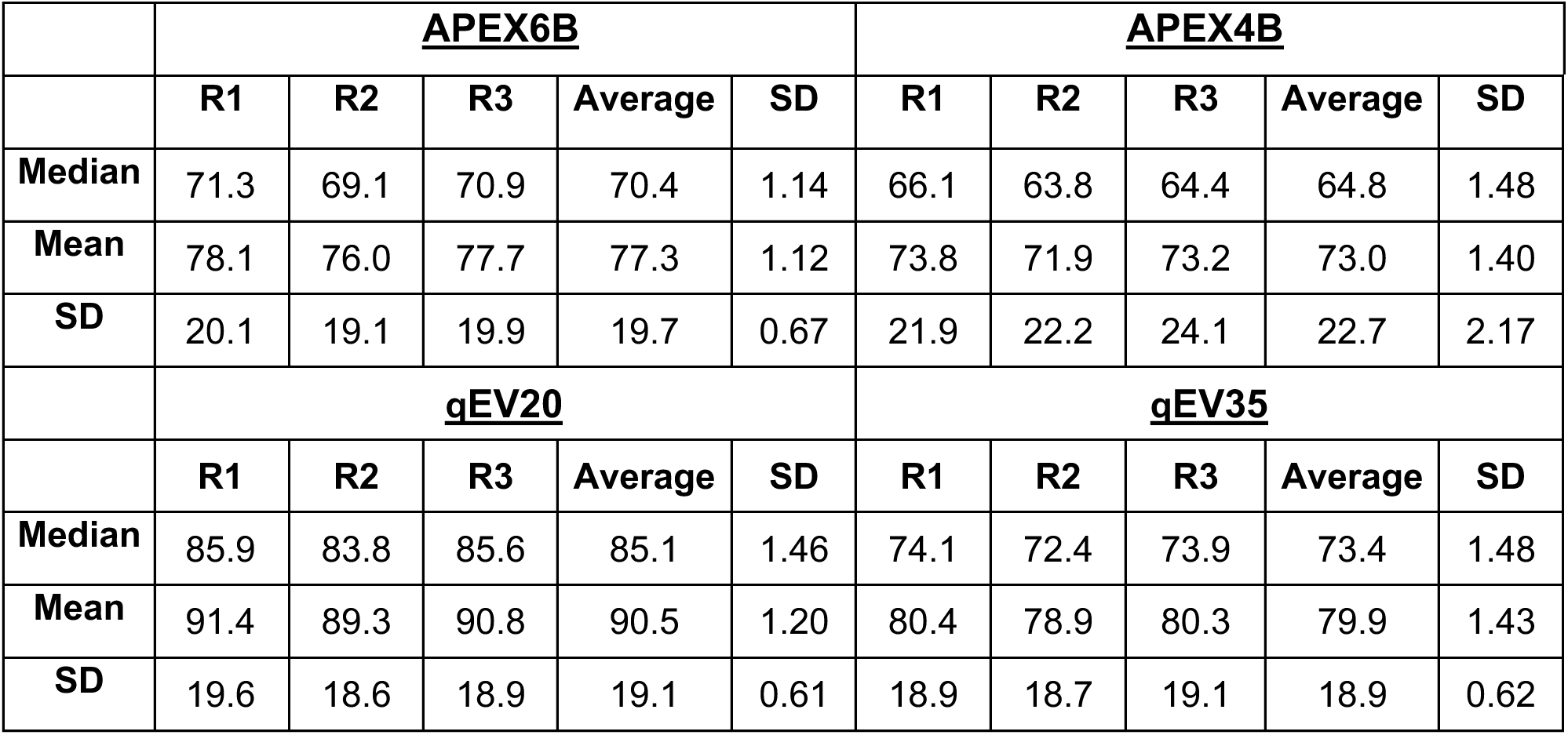
EndoC-ßH1-derived EV size measured by Nanoflow Cytometry (nm). R=replicate.

**Table 3:** EndoC-ßH1-derived EVs particle concentration measured by Nanoflow Cytometry represented as particles/mL. R=replicate.

|  | **<u>APEX6B</u>** |  |  |  |  | **<u>APEX4B</u>** |  |  |  |  |
| --- | --- | --- | --- | --- | --- | --- | --- | --- | --- | --- |
|  | **R1** | **R2** | **R3** | **Average** | **SD** | **R1** | **R2** | **R3** | **Average** | **SD** |
| **CD9** | 1.36×10<sup>10</sup> | 1.37×10<sup>10</sup> | 1.43×10<sup>10</sup> | 1.39×10<sup>10</sup> | 3.79×10<sup>8</sup> | 7.23×10<sup>9</sup> | 7.93×10<sup>9</sup> | 7.89×10<sup>9</sup> | 7.68×10<sup>9</sup> | 3.93×10<sup>8</sup> |
| **CD63** | 1.36×10<sup>10</sup> | 1.32×10<sup>10</sup> | 1.39×10<sup>10</sup> | 1.36×10<sup>10</sup> | 3.51×10<sup>8</sup> | 1.83×10<sup>10</sup> | 1.20×10<sup>10</sup> | 5.19×10<sup>9</sup> | 1.18×10<sup>10</sup> | 6.56×10<sup>9</sup> |
| **CD81** | 1.39×10<sup>10</sup> | 1.40×10<sup>10</sup> | 1.42×10<sup>10</sup> | 1.40×10<sup>10</sup> | 1.53×10<sup>8</sup> | 8.54×10<sup>9</sup> | 8.68×10<sup>9</sup> | 8.17×10<sup>9</sup> | 8.46×10<sup>9</sup> | 2.64×10<sup>8</sup> |
|  | **<u>qEV20</u>** |  |  |  |  | **<u>qEV35</u>** |  |  |  |  |
|  | **R1** | **R2** | **R3** | **Average** | **SD** | **R1** | **R2** | **R3** | **Average** | **SD** |
| **CD9** | 1.52×10<sup>10</sup> | 1.45×10<sup>10</sup> | 1.54×10<sup>10</sup> | 1.50×10<sup>10</sup> | 4.73×10<sup>8</sup> | 1.79×10<sup>10</sup> | 1.65×10<sup>10</sup> | 1.84×10<sup>10</sup> | 1.76×10<sup>10</sup> | 9.85×10<sup>8</sup> |
| **CD63** | 1.54×10<sup>10</sup> | 1.63×10<sup>10</sup> | 1.58×10<sup>10</sup> | 1.58×10<sup>10</sup> | 4.51×10<sup>8</sup> | 1.75×10<sup>10</sup> | 1.96×10<sup>10</sup> | 1.79×10<sup>10</sup> | 1.83×10<sup>10</sup> | 1.12×10<sup>9</sup> |
| **CD81** | 1.61×10<sup>10</sup> | 1.65×10<sup>10</sup> | 1.66×10<sup>10</sup> | 1.64×10<sup>10</sup> | 2.65×10<sup>8</sup> | 1.36×10<sup>10</sup> | 1.84×10<sup>10</sup> | 1.82×10<sup>10</sup> | 1.81×10<sup>10</sup> | 4.16×10<sup>8</sup> |

**Table 4:** EndoC-ßH1-derived EVs percentages of positive particles for the indicated tetraspanin marker measured by Nanoflow Cytometry. R=replicate.

|  | <u>**APEX6B**</u> |  |  |  |  | <u>**APEX4B**</u> |  |  |  |  |
| --- | --- | --- | --- | --- | --- | --- | --- | --- | --- | --- |
|  | **R1** | **R2** | **R3** | **Average** | **SD** | **R1** | **R2** | **R3** | **Average** | **SD** |
| **CD9** | 55.4% | 53.5% | 54.3% | 54.4% | 0.95% | 42.6% | 45% | 46.6% | 44.7% | 2.01% |
| **CD63** | 6.2% | 7.7% | 6.3% | 6.7% | 0.84% | 10.3% | 14.1% | 16.8% | 13.7% | 3.27% |
| **CD81** | 62.4% | 61.5% | 61.1% | 61.7% | 0.67% | 47.6% | 53.6% | 57.3% | 52.8% | 4.90% |
|  | <u>**qEV20**</u> |  |  |  |  | <u>**qEV35**</u> |  |  |  |  |
|  | **R1** | **R2** | **R3** | **Average** | **SD** | **R1** | **R2** | **R3** | **Average** | **SD** |
| **CD9** | 52.4% | 50.5% | 51.4% | 51.4% | 0.95% | 52.7% | 50.3% | 52.6% | 51.9% | 1.36% |
| **CD63** | 12.6% | 14.6% | 10.6% | 12.6% | 2.00% | 10.5% | 12.8% | 9.7% | 11.0% | 1.61% |
| **CD81** | 43.6% | 45.4% | 43.7% | 44.2% | 1.01% | 51.1% | 48.8% | 50.0% | 50.0% | 1.15% |

### Characterization of Human Plasma Isolated Fractions by Size Exclusion Chromatography

Human plasma EVs can be co-isolated with contaminants such as protein aggregates, lipoproteins, or viruses. To characterize the presence of some of these contaminants in the fractions isolated from human plasma using different size exclusion chromatography columns (Figure 3A), we detected the presence of human serum albumin (HSA) by Western blot (Figure 3B-D). HSA was primarily detected in F3-6 for the 20 nm columns, while in the 35 nm column, it was detected mostly in fractions F5-F6 (Figure 3B-D). Furthermore, the presence of another contaminant Apolipoprotein A (APOA), the primary protein component of high-density lipoprotein, was measured by ELISA. For the APEX columns, we found APOA mostly in F3-6, while IZON columns had high APOA content in all fractions (Figure 3E). When we measured particle concentration by tetraspanin ELISA, F3 had the highest concentration for both APEX columns - 3.65×10^10^±4.35×10^9^ particles/mL for the APEX6B column and 3.03×10^10^±4.30×10^9^ particles/mL for the APEX4B. F2 was the most particle-dense with 2.11×10^10^±3.52×10^9^ particles/mL for the qEV20 column and F1 for qEV35 column with 2.12×10^10^±6.20×10^9^ particles/mL (Figure 3F). In general, for all columns the fractions that usually are combined and concentrated to isolate EVs show the highest particle concentration. F5 and F6 have similar amounts of particles for both columns, although these are not included for downstream EV analysis.

**Figure 3:**
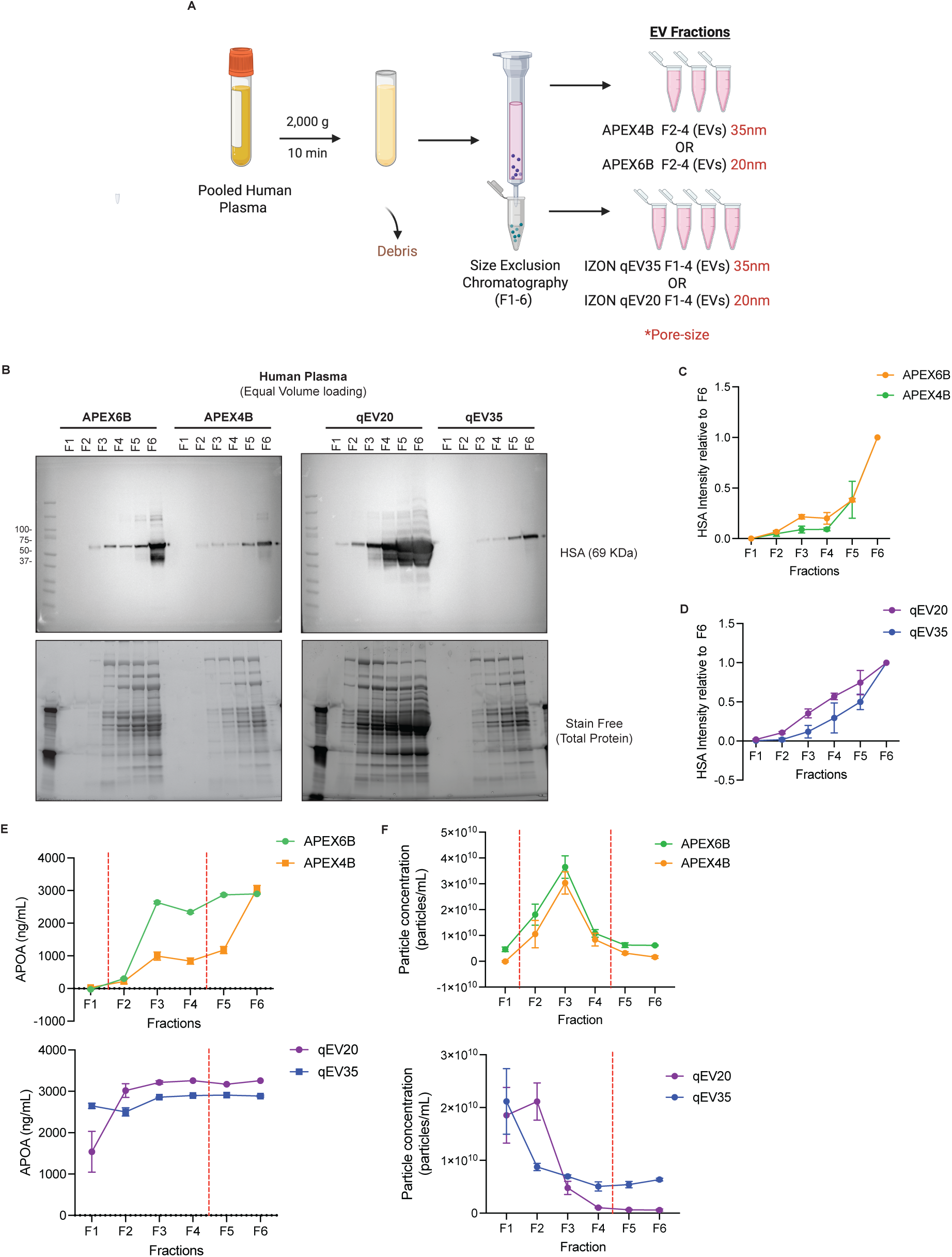
Comparison of isolated fractions from Human Plasma using different SEC columns. **A**. Schematic Illustration of Isolation workflow **B-D**. Representative Western blot and quantification of contaminant Human Serum Albumin (HSA) (*n*=3). **E.** APOA concentration in the isolated fractions quantified by an APOA ELISA (*n*=3) **F.** Particle concentration in the isolated fractions from 35 nm and 20 nm SEC columns quantified by TSPAN ELISA (*n*=3) (Data is shown as mean particles/ml ± SD). The red dotted line separates the fractions used for EV isolations, F2-F4 for APEX and F1-F4 for IZON.

### Comparison of Isolated EVs from Human Plasma

We characterized the morphology of human plasma-derived EVs using TEM. The EVs isolated with both columns, pooled from their respective EV fractions, displayed the typical cup-shaped morphology; however, EVs isolated using the APEX6B column appeared smaller in size (Figure 4A). We analyzed the size and concentration of the EVs by microfluidic resistive pulse sensing (MRPS). For both columns, the isolated EVs had a diameter of 60-80 nm as expected (Figure 4B). When analyzing the particle concentration, the APEX6B column yielded 8.83×10^12^±4.10×10^11^ particles/mL and the APEX4B column yielded 5.03×10^11^±3.85×10^11^ particles/mL. The qEV20 column yielded 1.96×10^11^±1.95×10^11^ particles/mL, while the qEV35 column yielded 6.00×10^10^±2.17×10^10^ particles/mL. When analyzed by nanoflow cytometry, mean EV diameter was 77.4±25.1 nm for the APEX6B column, 75.3±23.5 nm for APEX4B, 77.7±23.8 nm for qEV20 and 75.2±23.0 nm for qEV35 (Table 5). We used immunoblotting to check the presence of EV markers (Alix, CD63), and a negative control (GM130) (Figure 4C), as recommended by the MISEV guidelines. APEX6B EVs showed significantly higher CD63 expression than APEX4B EVs, but no significant differences in expression were observed for Alix or GM130 between any columns (Figure 4D). We calculated EV particle concentrations using a tetraspanin ELISA by taking the total concentration of the EV fractions for each column. Here, EVs isolated from APEX6B (2.18×10^10^±3.17×10^9^ particles/mL) were found to have significantly increased concentration to EVs isolated from APEX4B (1.64×10^10^±1.71×10^9^ particles/mL), but no difference in concentration was observed between EVs isolated from qEV20 (1.14×10^10^±4.86×10^8^ particles/mL) and qEV35 (1.05×10^10^±1.85×10^9^ particles/mL) (Figure 4E). Additionally, we also measured the total protein concentration of EV samples isolated from both columns using a BCA assay, where we found protein content to be significantly increased in 20 nm columns (Figure 4F). To characterize the relative sample purity, we also calculated the ratio of EV particle concentration to protein concentration as previously described^29^. We observed that APEX 4B samples exhibited a significantly higher particle/protein ratio than APEX6B samples, while no differences were observed between IZON columns (Figure 4G). Furthermore, we characterized the presence of contaminant APOA in the isolated EVs; APEX6B EVs contained more co-isolated APOA than APEX4B samples, whereas IZON columns showed no differences (Figure 4H).

**Figure 4:**
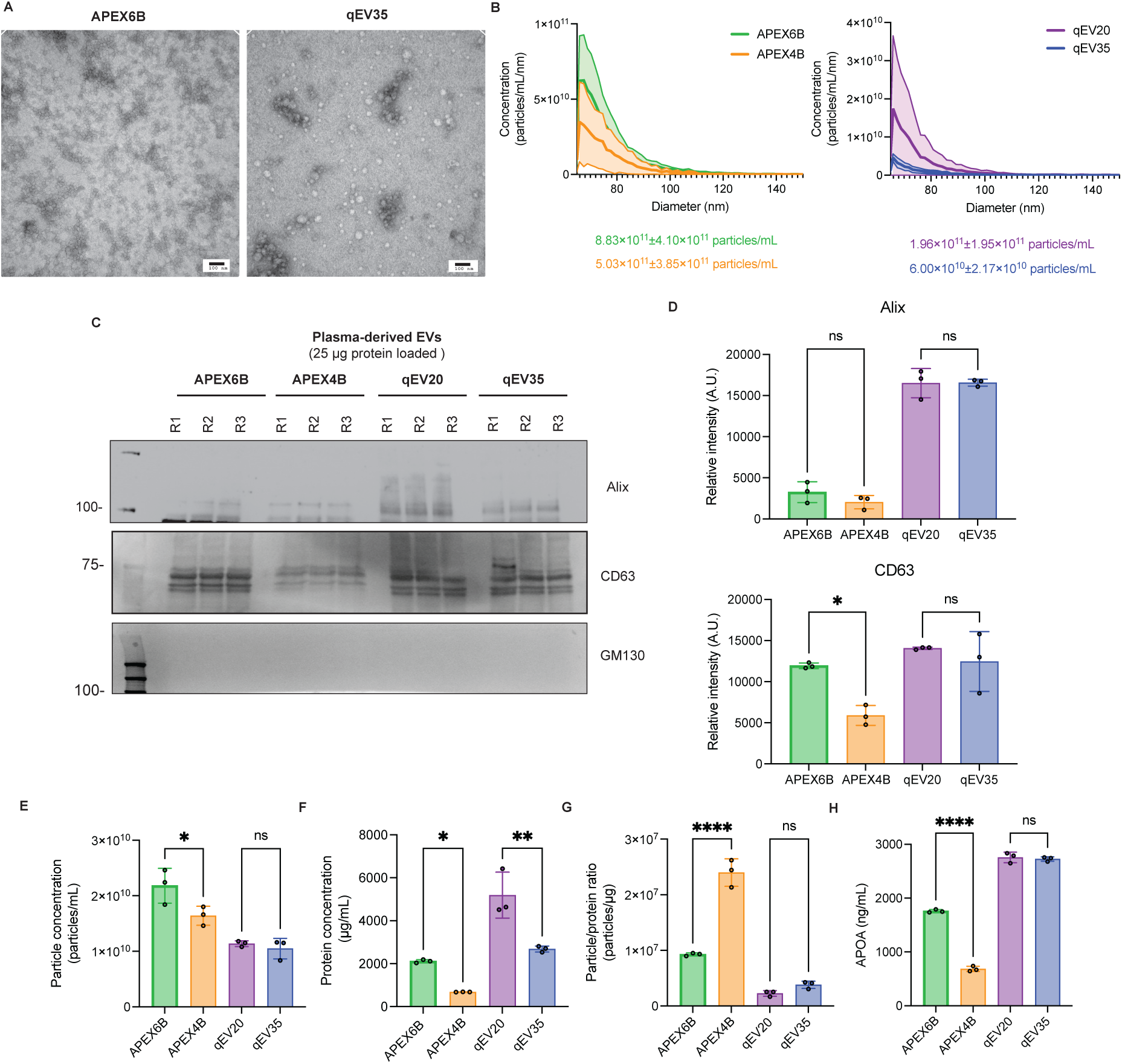
Characterization of EVs isolated from Human Plasma by different SEC columns. **A**. Transmission electron microscopy representative images, Scale=100nm. **B**. Size distribution measured by microfluidic resistive pulse sensing represented as mean±SD (*n*=3).**C-D**. Representative Western blot and quantification of EV markers of human plasma EVs (*n*=3). **E**. Human plasma EVs protein concentration by BCA (*n*=3). **F.** Particle concentration of the isolated EVs (*n*=3). **G**. Particle concentration to protein ratio of isolated EVs (*n*=3) **H**. APOA concentration in isolated EVs from SEC columns quantified by an APOA ELISA (*n*=3). All Data is represented as mean ± SD and *n*=3. ns=non-significant, \**p*<0.05, \*\**p*<0.01, \*\*\**p*≤0.001, \*\*\*\**p≤*0.0001. R=replicate.

**Table 5:** Human-pooled-plasma-derived EV size measured by Nanoflow Cytometry (nm). R=replicate.

|  | **<u>APEX6B</u>** |  |  |  |  | **<u>APEX4B</u>** |  |  |  |  |
| --- | --- | --- | --- | --- | --- | --- | --- | --- | --- | --- |
|  | **R1** | **R2** | **R3** | **Average** | **SD** | **R1** | **R2** | **R3** | **Average** | **SD** |
| **Median** | 69.8 | 70.8 | 70.2 | 70.3 | 0.50 | 69.2 | 69.2 | 68.2 | 68.9 | 0.58 |
| **Mean** | 77.5 | 77.5 | 77.1 | 77.4 | 0.23 | 75.5 | 75.7 | 74.8 | 75.3 | 0.47 |
| **SD** | 27.4 | 23.7 | 24.1 | 25.1 | 2.03 | 23.4 | 23.3 | 23.7 | 23.5 | 0.21 |
|  | **<u>qEV20</u>** |  |  |  |  | **<u>qEV35</u>** |  |  |  |  |
|  | **R1** | **R2** | **R3** | **Average** | **SD** | **R1** | **R2** | **R3** | **Average** | **SD** |
| **Median** | 70.8 | 70.2 | 70.8 | 70.6 | 0.35 | 68.8 | 68.8 | 68.2 | 68.6 | 0.35 |
| **Mean** | 77.9 | 77.3 | 77.9 | 77.7 | 0.35 | 75.3 | 75.9 | 74.5 | 75.2 | 0.70 |
| **SD** | 22.8 | 24.1 | 24.4 | 23.8 | 0.85 | 22.1 | 24.6 | 22.2 | 23.0 | 1.42 |

### Characterization of RNA cargo in Isolated EVs from EndoC-βH1 conditioned cell culture media and Human Plasma

Total RNA was isolated and characterized from EndoC-βH1-derived EVs and human plasma-derived EVs obtained by pooling the corresponding fractions from both columns. Total RNA quantification using a NanoDrop spectrophotometer from EndoC-βH1-derived EVs (Figure 5A) and human-plasma-derived EVs (Figure 5B) showed no significant differences between columns. EndoC-βH1-derived EV RNA profile showed enrichment in short RNAs (200-2000 nt) (Figure 5C and 5E-H), while human-plasma-derived EVs were mostly enriched in small RNAs (<200 nt) (Figure 5D and 5I-L) for columns of both pore sizes. To validate the use of different RNA species as biomarkers, we performed a qRT-PCR using microRNAs (miR). We characterized miR-21-5p which is known to be abundant in EndoC-βH1-derived EVs ^30^, as well as miR-30-d-5p expressed in pancreatic β-cells^31,32^ and human plasma EVs ^33^.No significant differences were observed in the expression of these miRNAs in EndoC-βH1-derived EVs (Figure 5M) or plasma-derived EVs (Figure 5H) isolated with the different columns.

**Figure 5:**
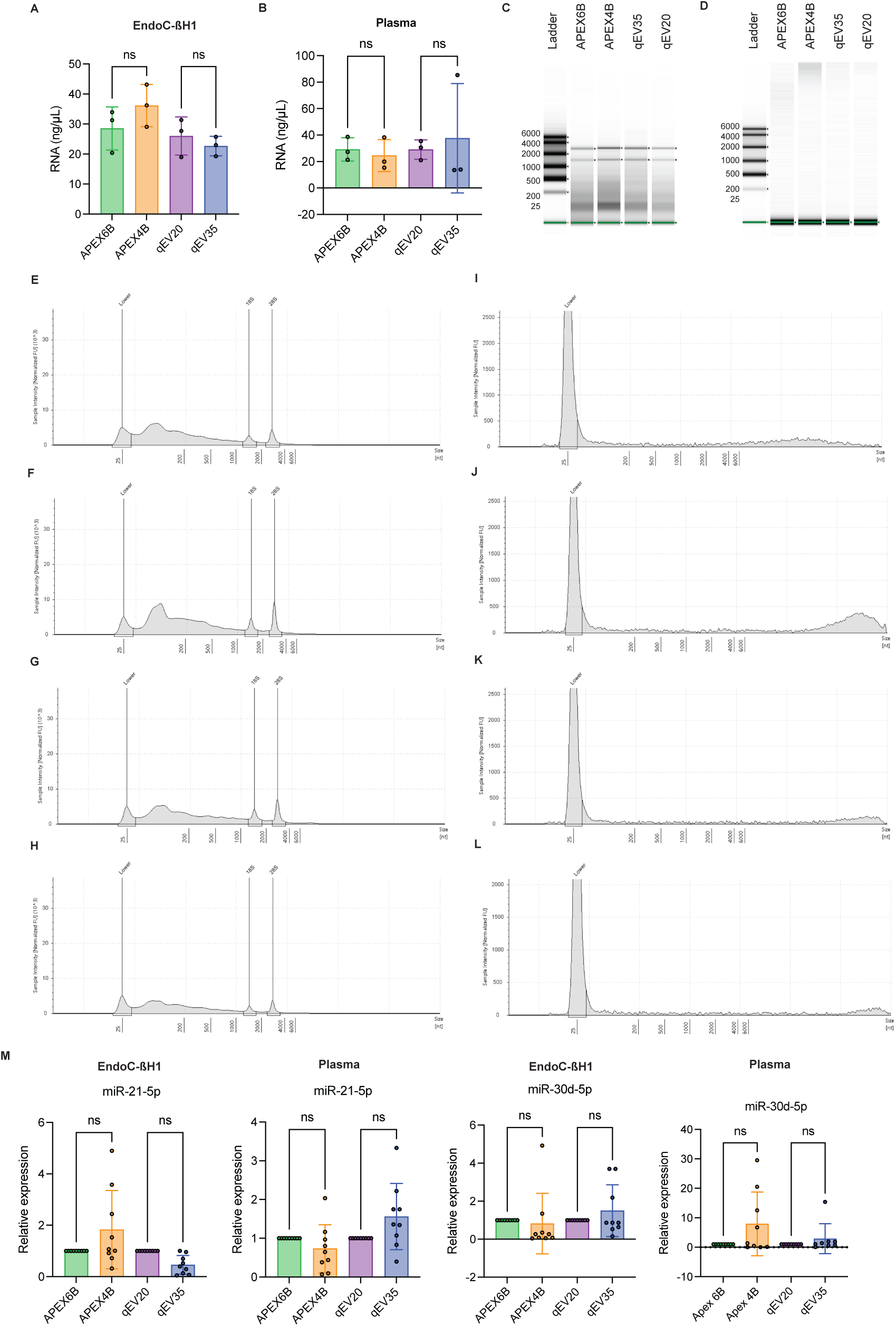
Characterization of nucleic acid cargo in EVs isolated by different SEC columns. **A.** EndoC-ßH1-derived EV RNA concentration measured by Nanodrop (*n*=3). **B.** Human plasma-derived EV derived RNA concentration measured by Nanodrop (*n*=3). **C-L.** EndoC-ßH1-derived EV RNA Densitometry traces profiles by TapeStation (*n*=3) **D.** Human plasma-derived EV RNA Densitometry traces profiles by TapeStation (*n*=3) **M.** miR-21-5p and miR-30d-5p expression normalized with cel-miR-39-3p (*n*=3). Student’s t-test non-significant (ns).

## Discussion

Extracellular vesicles (EVs) can be isolated using methods that differ in EV yields and purities^16,34,35^. One of the most widely used methods for EV isolation is size-exclusion chromatography (SEC), which is preferred for its rapidness and reproducibility^36^. However, commercially available SEC columns present different pore sizes, which may influence not only EV recovery but also the removal of other extracellular particles. The presence of these ‘contaminating co-isolates’ can significantly affect downstream analyses, such as analyses of EV cargo^9,11,12^. In this study, we compared SEC columns with different pore sizes - 20 nm (APEX6B and qEV20) and 35 nm (APEX4B and qEV35) - from two manufacturers, in the context of pancreatic biology, focusing on their applicability for biomarker discovery in Type 1 Diabetes.

Our results show differences in the presence of extracellular co-isolates based on the pore size and fractions used for EV isolation. In a western blot of EndoC-βH1 conditioned media, the presence of bovine serum, a known contaminant that may affect the purity of EV isolations^9,10^, was mostly found in all fractions for the 20 nm pore size columns and in the later fractions for the 35 nm size columns. These findings suggest that the 35 nm pore size column elutes contaminants in later fractions not used for EV isolation, potentially leading to a cleaner isolation of EVs from the earlier fractions, although restricting the fractions for EV isolations may also serve to decrease any contaminating free protein. This trend was also seen in plasma derived EVs, where the removal of major plasma contaminants such as HSA was more efficient with the 35 nm pore size columns. Samples isolated using 35 nm pore size columns contained practically to no HSA in F2-4, whereas isolations with 20 nm size pore columns had detectable HSA in F3-F6. This difference may explain the lower yield observed since the presence of albumin might be interfering with the EV yield and purity as previously shown^20^. Overall, these results show a general decrease in plasma contaminants on the EV-containing fractions isolated by 35 nm columns compared to 20 nm columns, consistent with previous studies showing that smaller pore sizes in SEC columns increase EV yield as well as contaminants^37,38^.

Prior literature has suggested that it is insufficient or inadequate to characterize EV quantity only based on nanoparticle tracking or protein content, particularly for plasma derived EVs, as contaminants are likely to affect these measurements^20,29^. TEM confirmed that EVs from both columns maintained the expected cup-shaped morphology. When characterized by microfluidic resistive pulse sensing (MRPS) and nanoflow cytometry we observed larger size distribution for APEX6B and qEV20 (20 nm columns) both columns. However, particle concentration measured by a tetraspanin ELISA showed a higher particle concentration in F3-4 for APEX6B and APEX4B, and F1-2 for qEV35 and qEV20. When the corresponding EV-containing fractions were pooled (F2-4 for APEX6B and APEX4B, F1-4 for qEV35 and qEV20), the columns with 20 nm pore size consistently yielded higher particle concentrations. Interestingly, although 20 nm pore size columns showed a higher overall protein concentration in EV isolates from human plasma, it did not translate to higher EV content. However, when the particle-to-protein ratio was measured, APEX4B and qEV35 (both 35 nm columns) showed a higher value, indicating both greater yield and higher purity for both EndoC-βH1 and plasma-derived EVs.

In addition to measuring EV yield and purity, we conducted an analysis of EV cargo and how it may differ between the two column pore sizes. We measured by Western blot the relative quantity of cargo, membrane, and cell contaminant protein markers (Alix, CD63, GM130, respectively)^28^ for EVs derived from conditioned media and human plasma. Although there were differences in expression for some EV markers, there was no general trend suggesting EVs isolated from either column contain greater overall EV marker expression. We also performed nanoflow cytometry with EVs derived from EndoC-ßH1 conditioned media to check for expression of the tetraspanin surface EV markers CD9, CD63, and CD81. All three of these proteins were shown to be expressed at similar levels for EVs isolated from all columns. These findings suggest that EV protein cargo seems to be conserved between the two pore sizes of the size-exclusion columns, despite the previously mentioned above differences in EV recovery and contaminant removal.

Another critical observation was the RNA cargo of EVs isolated by both methods. Total RNA quantification and RNA integrity analyses revealed no major differences in RNA yield or profiles between the columns, suggesting that RNA cargo preservation is not compromised by pore size differences. These results suggest that both columns are suitable for RNA-based downstream applications, although 20 nm pore size may offer slight advantages in EV recovery consistency and 35 nm pore size in contaminant removal that might affect other downstream RNA analysis, at least for the types of samples tested here. Nonetheless, with careful attention to the use of appropriate fractions with the minimal amount of contaminating co-isolates, it would be reasonable to use either method for EV cargo analyses, especially if a particular investigator has easier access of familiarity with one particular type of platform.

In summary, this study highlights that SEC column pore size is a key determinant in optimizing EV isolation from both conditioned media and human plasma. The 20 nm size pore column consistently demonstrated higher EV yields and 35 nm size pore columns lower contaminant levels. This is particularly significant in cell types with lower EV secretion such as EndoC-βH1 cells, where higher yield becomes a critical factor. Importantly, the 20 nm pore size columns can achieve this without compromising purity, since the presence of some contaminants might not be compatible with some downstream applications. An important consideration when isolating EVs for biomarker discovery in diseases such as Type 1 Diabetes are limiting factors such as sample availability and biological variability. In conclusion, overall, these findings support the use of smaller pore size SEC columns for applications requiring high-purity EVs, particularly in settings with challenging sample compositions or where downstream biomarker analysis is sensitive to contaminant interference.

## Acknowledgments

This work was funded by the National Institute of Diabetes and Digestive and Kidney Diseases under grant number 5R01DK133847 and NIH U01 DK135095 (R.N.K and S.D). The authors would like to thank Das lab members for their feedback.

## References

1. Silva, T. F. et al. Extracellular vesicle heterogeneity through the lens of multiomics. Cell Reports Medicine 6, 102161 (2025).

2. van Niel, G., D’Angelo, G. & Raposo, G. Shedding light on the cell biology of extracellular vesicles. Nat Rev Mol Cell Biol 19, 213–228 (2018).

3. Hinestrosa, J. P. et al. Early-stage multi-cancer detection using an extracellular vesicle protein-based blood test. Commun Med 2, 1–9 (2022).

4. Blaser, M. C. et al. Multiomics of Tissue Extracellular Vesicles Identifies Unique Modulators of Atherosclerosis and Calcific Aortic Valve Stenosis. Circulation 148, 661–678 (2023).

5. Kerr, N. et al. Inflammasome Proteins in Serum and Serum-Derived Extracellular Vesicles as Biomarkers of Stroke. Front Mol Neurosci 11, 309 (2018).

6. Garcia-Contreras, M. et al. Plasma-derived exosome characterization reveals a distinct microRNA signature in long duration Type 1 diabetes. Sci Rep 7, 5998 (2017).

7. Kalluri, R. & LeBleu, V. S. The biology, function, and biomedical applications of exosomes. Science 367, eaau6977 (2020).

8. Cheng, L. & Hill, A. F. Therapeutically harnessing extracellular vesicles. Nat Rev Drug Discov 21, 379–399 (2022).

9. Lehrich, B. M., Liang, Y. & Fiandaca, M. S. Foetal bovine serum influence on in vitro extracellular vesicle analyses. J Extracell Vesicles 10, e12061 (2021).

10. Urzì, O., Olofsson Bagge, R. & Crescitelli, R. The dark side of foetal bovine serum in extracellular vesicle studies. J Extracell Vesicles 11, e12271 (2022).

11. Pham, C. V. et al. Bovine extracellular vesicles contaminate human extracellular vesicles produced in cell culture conditioned medium when ‘exosome-depleted serum’ is utilised. Archives of Biochemistry and Biophysics 708, 108963 (2021).

12. Shelke, G. V., Lässer, C., Gho, Y. S. & Lötvall, J. Importance of exosome depletion protocols to eliminate functional and RNA-containing extracellular vesicles from fetal bovine serum. J Extracell Vesicles 3, (2014).

13. Sódar, B. W. et al. Low-density lipoprotein mimics blood plasma-derived exosomes and microvesicles during isolation and detection. Sci Rep 6, 24316 (2016).

14. Stolk, M. & Seifert, M. Protein contaminations impact quantification and functional analysis of extracellular vesicle preparations from mesenchymal stromal cells. J Stem Cells Regen Med 11, 44–47 (2015).

15. Cocozza, F., Grisard, E., Martin-Jaular, L., Mathieu, M. & Théry, C. SnapShot: Extracellular Vesicles. Cell 182, 262–262.e1 (2020).

16. Suresh, P. S. & Zhang, Q. Comprehensive Comparison of Methods for Isolation of Extracellular Vesicles from Human Plasma. J Proteome Res 24, 2956–2967 (2025).

17. Saludas, L. et al. Isolation methods of large and small extracellular vesicles derived from cardiovascular progenitors: A comparative study. European Journal of Pharmaceutics and Biopharmaceutics 170, 187–196 (2022).

18. Ter-Ovanesyan, D. et al. Improved isolation of extracellular vesicles by removal of both free proteins and lipoproteins. eLife 12, e86394 (2023).

19. Takov, K., Yellon, D. M. & Davidson, S. M. Comparison of small extracellular vesicles isolated from plasma by ultracentrifugation or size-exclusion chromatography: yield, purity and functional potential. J Extracell Vesicles 8, 1560809 (2019).

20. Brennan, K. et al. A comparison of methods for the isolation and separation of extracellular vesicles from protein and lipid particles in human serum. Sci Rep 10, 1039 (2020).

21. Chou, C.-Y. et al. Improving the Purity of Extracellular Vesicles by Removal of Lipoproteins from Size Exclusion Chromatography- and Ultracentrifugation-Processed Samples Using Glycosaminoglycan-Functionalized Magnetic Beads. ACS Appl. Mater. Interfaces 16, 44386–44398 (2024).

22. Tsonkova, V. G. et al. The EndoC-βH1 cell line is a valid model of human beta cells and applicable for screenings to identify novel drug target candidates. Mol Metab 8, 144–157 (2017).

23. Hastoy, B. et al. Electrophysiological properties of human beta-cell lines EndoC-βH1 and -βH2 conform with human beta-cells. Sci Rep 8, 16994 (2018).

24. Ryaboshapkina, M. et al. Characterization of the Secretome, Transcriptome, and Proteome of Human β Cell Line EndoC-βH1. Molecular & Cellular Proteomics 21, (2022).

25. Ravassard, P. et al. A genetically engineered human pancreatic β cell line exhibiting glucose-inducible insulin secretion. J Clin Invest 121, 3589–3597 (2011).

26. Basile, G. et al. Excess pancreatic elastase alters acinar-β cell communication by impairing the mechano-signaling and the PAR2 pathways. Cell Metabolism 35, 1242–1260.e9 (2023).

27. De Jesus, D. F. et al. m6A mRNA Methylation Regulates Human β-Cell Biology in Physiological States and in Type 2 Diabetes. Nat Metab 1, 765–774 (2019).

28. Welsh, J. A. et al. Minimal information for studies of extracellular vesicles (MISEV2023): From basic to advanced approaches. J Extracell Vesicles 13, e12404 (2024).

29. Webber, J. & Clayton, A. How pure are your vesicles? Journal of Extracellular Vesicles 2, 19861 (2013).

30. Lakhter, A. J. et al. Beta cell extracellular vesicle miR-21-5p cargo is increased in response to inflammatory cytokines and serves as a biomarker of type 1 diabetes. Diabetologia 61, 1124–1134 (2018).

31. Tang, X., Muniappan, L., Tang, G. & Ozcan, S. Identification of glucose-regulated miRNAs from pancreatic {beta} cells reveals a role for miR-30d in insulin transcription. RNA 15, 287–293 (2009).

32. Mao, Y. et al. Transgenic overexpression of microRNA-30d in pancreatic beta-cells progressively regulates beta-cell function and identity. Sci Rep 12, 11969 (2022).

33. Frørup, C. et al. Plasma Exosome-Enriched Extracellular Vesicles From Lactating Mothers With Type 1 Diabetes Contain Aberrant Levels of miRNAs During the Postpartum Period. Front Immunol 12, 744509 (2021).

34. Wallis, R. et al. Isolation methodology is essential to the evaluation of the extracellular vesicle component of the senescence-associated secretory phenotype. J Extracell Vesicles 10, e12041 (2021).

35. Robinson, S. D. et al. Confirming size-exclusion chromatography as a clinically relevant extracellular vesicles separation method from 1mL plasma through a comprehensive comparison of methods. BMC Methods 1, 7 (2024).

36. Corso, G. et al. Reproducible and scalable purification of extracellular vesicles using combined bind-elute and size exclusion chromatography. Sci Rep 7, 11561 (2017).

37. Bracht, J. W. P. et al. Choice of size-exclusion chromatography column affects recovery, purity, and miRNA cargo analysis of extracellular vesicles from human plasma. Extracell Vesicles Circ Nucl Acids 5, 497–508 (2024).

38. György, B. et al. Effect of the 35 nm and 70 nm Size Exclusion Chromatography (SEC) Column and Plasma Storage Time on Separated Extracellular Vesicles. Current Issues in Molecular Biology 46, 4337–4357 (2024).

